# The *Streptococcus mutans* Rhamnose-glucose Polysaccharide Plays an Important Role in Oxidative Stress Resistance and Iron Homeostasis

**DOI:** 10.1101/2022.11.28.518218

**Authors:** Andrew P. Bischer, Roberta C. Faustoferri, Christopher J. Kovacs, Steven R. Gill, Robert G. Quivey

## Abstract

The rhamnose-glucose polysaccharide (RGP) of *Streptococcus mutans* is known to play an essential role in cell division and biofilm formation. It has become increasingly apparent that the structure plays a role in protection against external stress, including an undefined role in oxidative stress. In this study, we demonstrate that the loss of RGP impairs the uptake of glucose and the susceptibility of *S. mutans* to lethal concentrations of iron, while at the same time, generates elevated content of intracellular iron, a prominent producer of reactive oxygen species. Intracellular iron levels were significantly higher in a strain lacking rhamnose production (Δ*rmlD*), compared to a strain deficient in components of the RGP production pathway (Δ*rgpG*). The seemingly paradoxical discrepancy between iron uptake and iron susceptibility may be partially explained by increased expression of *dpr* and *sod*, genes that protect against oxidative stress and prime *S. mutans* against oxidative stressors such as H_2_O_2_ and superoxide. Analysis of the promoter of the rhamnose biosynthesis gene *rmlD* indicated the presence of regulatory motifs, including Rex and PerR, which we have shown control expression for the genes responsible for rhamnose biosynthesis and RGP production. These results provide initial insights into the role that RGP plays in toleration of oxidative stress and suggest a complexity in regulation of the genes responsible for production of the RGP.

**Importance:** Dental caries, a disease caused by *Streptococcus mutans*, imposes significant healthcare burdens and productivity losses in the US. The outer structure of *S. mutans*, which is primarily composed of the rhamnose-glucose polysaccharide (RGP), is understudied, though it plays a crucial role in survival of the organism in response to environmental stress. Elucidating the protective role of RGP and the regulatory mechanisms that control the synthesis of RGP in *S. mutans* will lead to insights into the structure and function of cell wall polysaccharides in other streptococci. This study reveals a role for L-rhamnose, the building block of RGP, in mediating homeostasis of iron uptake, while the loss of rhamnose biosynthesis causes alterations in transcription of *sod* and *dpr*.

## Introduction

Streptococci are unlike many other Gram-positive bacteria in that they lack wall-teichoic acid structures, with their cell walls decorated by rhamnose-containing polysaccharides (1). This structure in the oral pathogen *Streptococcus mutans*, termed the rhamnose-glucose polysaccharide, or RGP (2), has recently come into focus due to studies that have demonstrated important functions for the RGP structure in cell division and stress protection (3–6). The first step in RGP construction is catalyzed by the enzyme RgpG, which adds an N-acetylglucosamine (GlcNAc) to an undecaprenol lipid carrier (7). After addition of the GlcNAc base, the rhamnan backbone of the structure is formed by the RgpA-catalyzed addition of the first L-rhamnose subunit (8), followed by the addition of subsequent rhamnose subunits and elongation of the backbone by RgpB and RgpF (8). The process culminates in the addition of the glucose side chain (RgpE, RgpH, and RgpI) (8, 9) and a glycerol phosphate modification to that side chain by Orf7 (6). The mature RGP is then exported to the exterior of the cell and crosslinked to the peptidoglycan (10). The RGP has been shown to play a protective role against environmental stressors, to coordinate cell wall synthesis proteins for division, and be required for proper cell morphology (3–6). In this work, we demonstrate, for the first time, how the RGP structure protects against stress from oxidative and acidic sources.

Oxidative stress is a major stressor encountered by *S. mutans* in the oral cavity. Due to the nature of constant inhalation, the oral cavity is an oxygen-rich location. Many commensal bacteria that reside in the mouth are known to produce large amounts of reactive oxygen species (ROS) (11). *Streptococcus gordonii* and *Streptococcus sanguinis* are common peroxigenic oral commensals, and major competitors of *S. mutans*, via their production of large quantities of hydrogen peroxide (12, 13). The constant bombardment of oxygen and hydrogen peroxide by these organisms allows for their reduction to more deleterious ROS such as superoxide anions and hydroxyl radicals (14).

In addition to hydrogen peroxide, metals, such as iron, have a well-characterized role in the formation of ROS and oxidative stress in *S. mutans* (15). Salivary iron levels have been measured between 0.1 μM and 1.0 μM (16), although the amount and availability of iron in the saliva very likely increases temporarily after the consumption of food. A classic example of this metal-mediated generation of ROS is the Fenton reaction, where ferrous iron reduces hydrogen peroxide into the highly-reactive hydroxyl radical (17). Production of these hydroxyl radicals, which can damage lipids, DNA, and amino acids, can have significant consequences on the cells (18).

*S. mutans* is not defenseless in the face of oxidative stressors. Like other streptococci, *S. mutans* does not produce catalase to reduce hydrogen peroxide (19), but rather the organism is protected from hydrogen peroxide-mediated ROS by the production of an alkyl hydroperoxide reductase (AhpCF) (20) that degrades hydrogen peroxide to water. To prevent the Fenton reaction, *S. mutans* produces Dpr to sequester intracellular iron (21). Dpr has been shown to be important for protection against reactive oxygen species and participates in competition with peroxigenic streptococci by limiting damage caused by H_2_O_2_ (22, 23). Superoxide dismutase has also been shown to play a role in the arms race against competing bacteria (23) by reducing superoxide anions (24). The NADH oxidase, NoxA, catalyzes the reduction of oxygen to water via the addition of electrons, allowing for a reduction in intracellular oxygen and the recycling of NADH (25, 26). These enzymes play a major role in protection of *S. mutans* against ROS, but are not the only enzymes involved in tolerating the presence of oxygen and hydrogen peroxide in the oral cavity.

The transcription factors PerR and Rex both modify transcription of genes that act in response to oxidative stress. PerR is a member of the Fur family of transcriptional regulators that regulate iron homeostasis and expression of oxidative stress response genes in streptococci (27–30). The genes *dpr* (SMU.540)*, ahpCF* (SMU.764-765), and *sod* (SMU.629) have all been shown to be negatively regulated, or repressed, by PerR (23, 28, 31, 32). Loss of PerR in *S. mutans* has been shown to result in decreased susceptibility to hydrogen peroxide (23). Rex, in contrast to PerR, can act as a transcriptional activator in *S. mutans*, and the loss of Rex has been demonstrated to cause elevated susceptibility to oxidative stress (33–35). Rex has also been shown to regulate expression of *noxA* (SMU.1117c), which encodes a NADH oxidase responsible for maintaining intracellular NADH/NAD^+^ levels (26, 33, 36).

While the number of studies directed at the RGP of *S. mutans* has grown in the last few years, there is still much about the structure that is not known, including the mechanism by which the polysaccharide protects the organism from stress. In this study, we show that the loss of the RGP reduces the organism’s susceptibility to oxidative stress-inducing iron. Unexpectedly, the loss of the RGP also resulted in increased intracellular levels of iron. We also demonstrate that glucose import and proton permeability are disrupted by RGP elimination, further highlighting dysfunction in transport and membrane stability in the RGP-deficient mutant strains. As further evidence of the role of the structure in protection against oxidative stress, we show that the operon responsible for RGP biosynthesis is regulated by the transcription factors Rex and PerR, both involved in modulating the oxidative stress response and iron homeostasis (28, 29, 34–38). Finally, we show that mutant strains lacking genes responsible for synthesis and assembly of RGP machinery likely overcome increased intracellular iron levels by upregulating expression of genes that are a part of the oxidative stress response pathways.

## Results

### *S. mutans* displays reduced susceptibility to iron-mediated killing in the absence of RGP

Previous research demonstrated that alteration, or total loss, of the RGP had significant effects on the ability of *S. mutans* to withstand oxidative stress (3, 4, 39). By extension, we wanted to determine if the RGP participated in protection against iron-mediated damage, which is generally through ROS production. First, we measured the growth rate of *S. mutans* strains deficient in an aspect of the RGP structure when grown in medium containing subinhibitory concentrations of iron, under both aerobic (Figure 1A) and microaerobic conditions (Figure 1B). Interestingly, the presence of iron in the culture medium appeared to cause no significant defects on the growth of any of the strains tested. The Δ*rgpF* (SMU.830), Δ*rgpG* (SMU.246), and Δ*rmlD* (SMU.824) strains all have been shown to be more susceptible to oxidative stress, but none of these strains displayed altered growth rates when exposed to sublethal levels of ferrous iron (3, 4, 39) in an aerobic or microaerobic environment (Table). Here, no significant changes were observed in the doubling time of any of the strains tested when grown in microaerobic conditions. In aerobic growth conditions, we found some interesting results, with improved growth among some strains rather than impaired growth. Again, the *rgpG* and *rmlD* deletion strains displayed no significant changes in doubling time, compared to growth in media without additional iron; but, the UA159, *rmlD*^+^, Δ*rgpF*, and Δ*orf7* (SMU.831) strains exhibited a significant decrease in doubling time, compared to growth in media without iron (Table 1). The *orf7* gene had previously been shown to encode a protein responsible for attachment of a glycerol phosphate moiety to the rhamnose chain, and was reported to be protective against zinc-mediated damage (6). Our data shows that, at least at subinhibitory concentrations of iron, the *orf7* deletion strain exhibited no sensitivity to iron-mediated stress (Figure 1A and 1B, Table 1).

**Figure 1.**
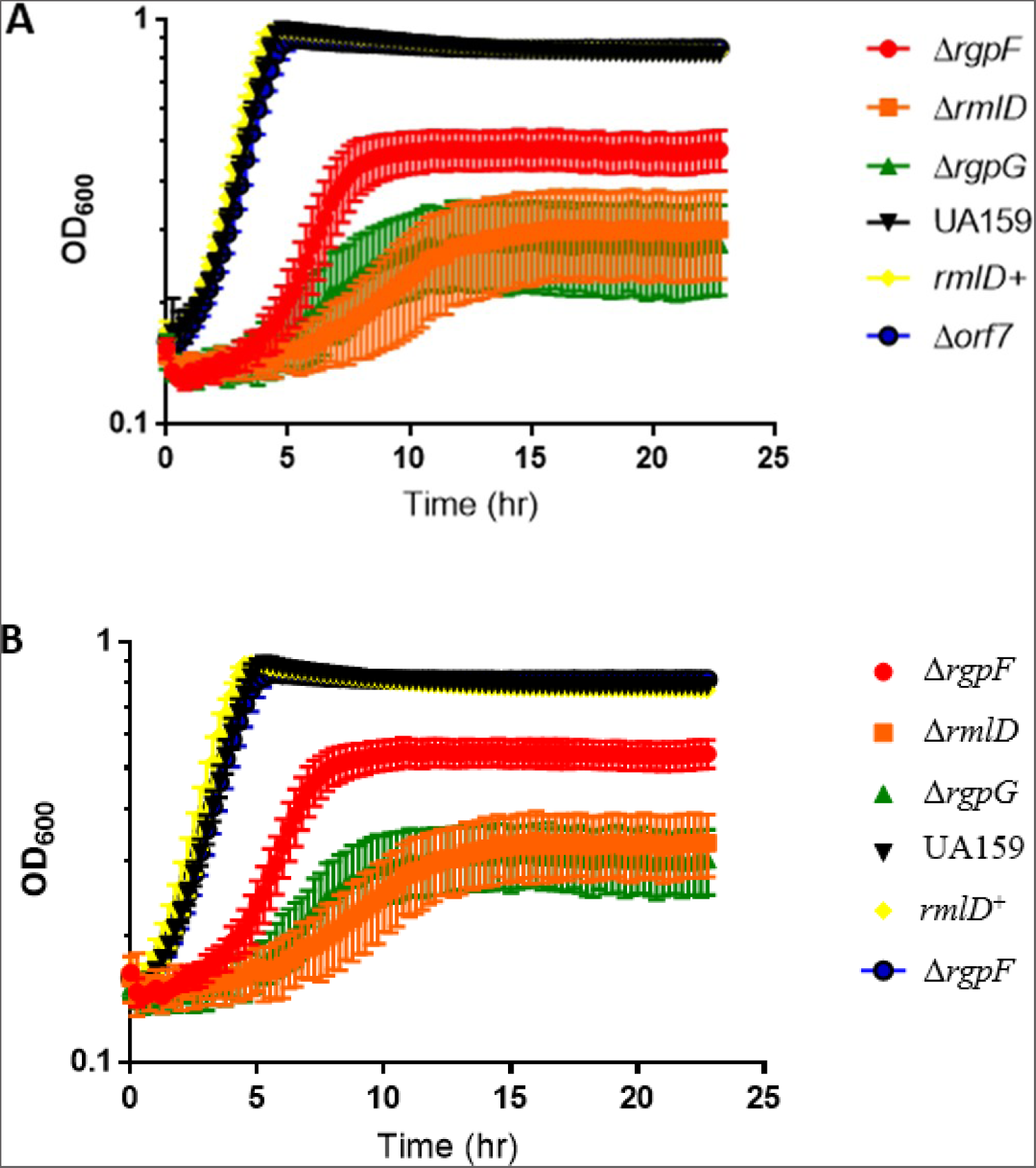
Loss or alteration of the *S. mutans* RGP does not affect growth of the organism in sub-lethal concentrations of ferrous iron. Strains of *S. mutans* carrying deletions in *rgpF*, *rmlD*, *rgpG*, Orf7, in addition to the parent strain, UA159, and the *rmlD* complement strain (*rmlD^+^*) were grown in BHI medium containing 1 mM ferrous ammonium sulfate in the absence of (Panel A) or presence of (Panel B) an oil overlay to create a microaerobic environment. Optical density (OD_600_) readings were taken at 15 minute intervals in a Bioscreen C Plate Reader (± standard deviation). Doubling times were calculated and listed in the table below each graph. Experiments were performed from 3 independent cultures of each strain, with 10 technical replicates per strain. Doubling times can be found in Table 1.

**Table 1:**
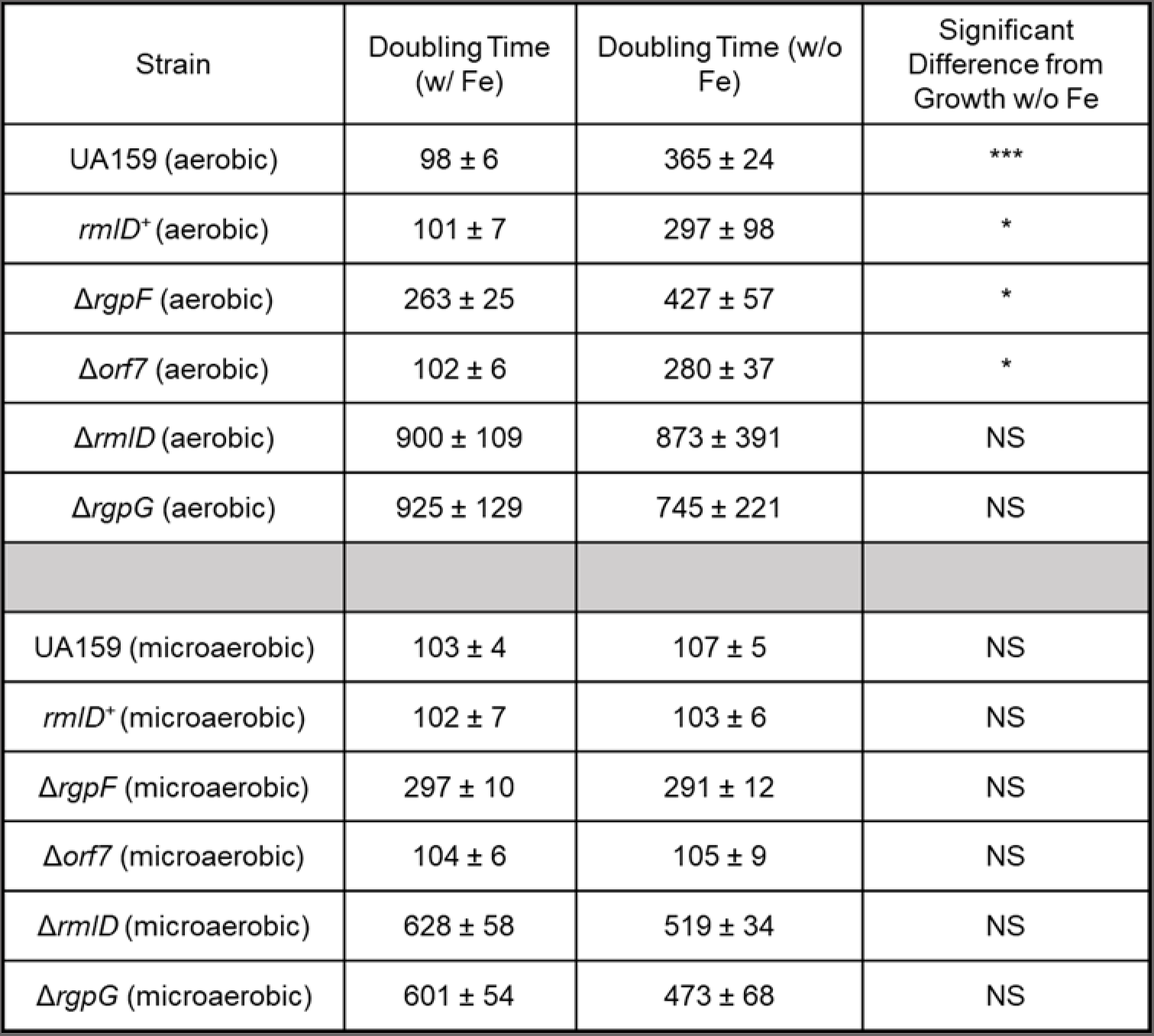
Doubling times of *S. mutans* strains displayed in Figure 1A & 1B. Aerobic indicates growth without the oil overlay (Fig. 1A) and microaerobic indicates growth with the oil overlay (Fig. 1B). Doubling times were determined as described in the Materials and Methods. Doubling times for strains grown without iron were previously determined (9, 38). Statistical significance was determined via pairwise comparison using a Student’s *t-*test between doubling time with iron and without iron. * indicates *p* < 0.05; **, *p* < 0.01; ***, *p* < 0.001.

Next, we used a survival assay to measure the response of the test strains to a lethal dose of iron, a ten-fold increased concentration compared to the concentration used in the growth experiments. We hypothesized that the mutant strains with an incomplete or ablated RGP would be more susceptible to this elevated level of iron due to their defects in stress tolerance. Surprisingly, we observed the opposite result. The strains completely devoid of the RGP, via deletion of *rmlD* or *rgpG*, displayed increased tolerance to the iron-mediated stress, compared to the UA159 strain (Figure 2). This finding was notable because in every other case where a stress agent was tested, strains with a defect in the RGP were typically more susceptible (3–5). The Δ*orf7* strain was again as equally susceptible as UA159, indicating that the glycerol phosphate may not be involved in protection against iron-mediated damage. Another interesting discovery from this experiment was the susceptibility of the Δ*rgpF* strain. RgpF is one of the glycosyltransferases required for extension of the rhamnose backbone of the RGP (4). This deletion strain exhibited an intermediate response to iron killing, compared to the other test strains, suggesting that perhaps the overall amount of rhamnose in the cell wall is linked to the phenomenon we observed. We repeated this assay, this time using lethal levels of H_2_O_2_, to determine if the results could be replicated in the presence of another oxidative stressor. We found that the Δ*rgpG* and Δ*rmlD* strains were both significantly more susceptible to H_2_O_2_, compared to UA159, confirming previous studies of susceptibility to oxidative stress (3, 4, 39) (Figure S1).

**Figure 2.**
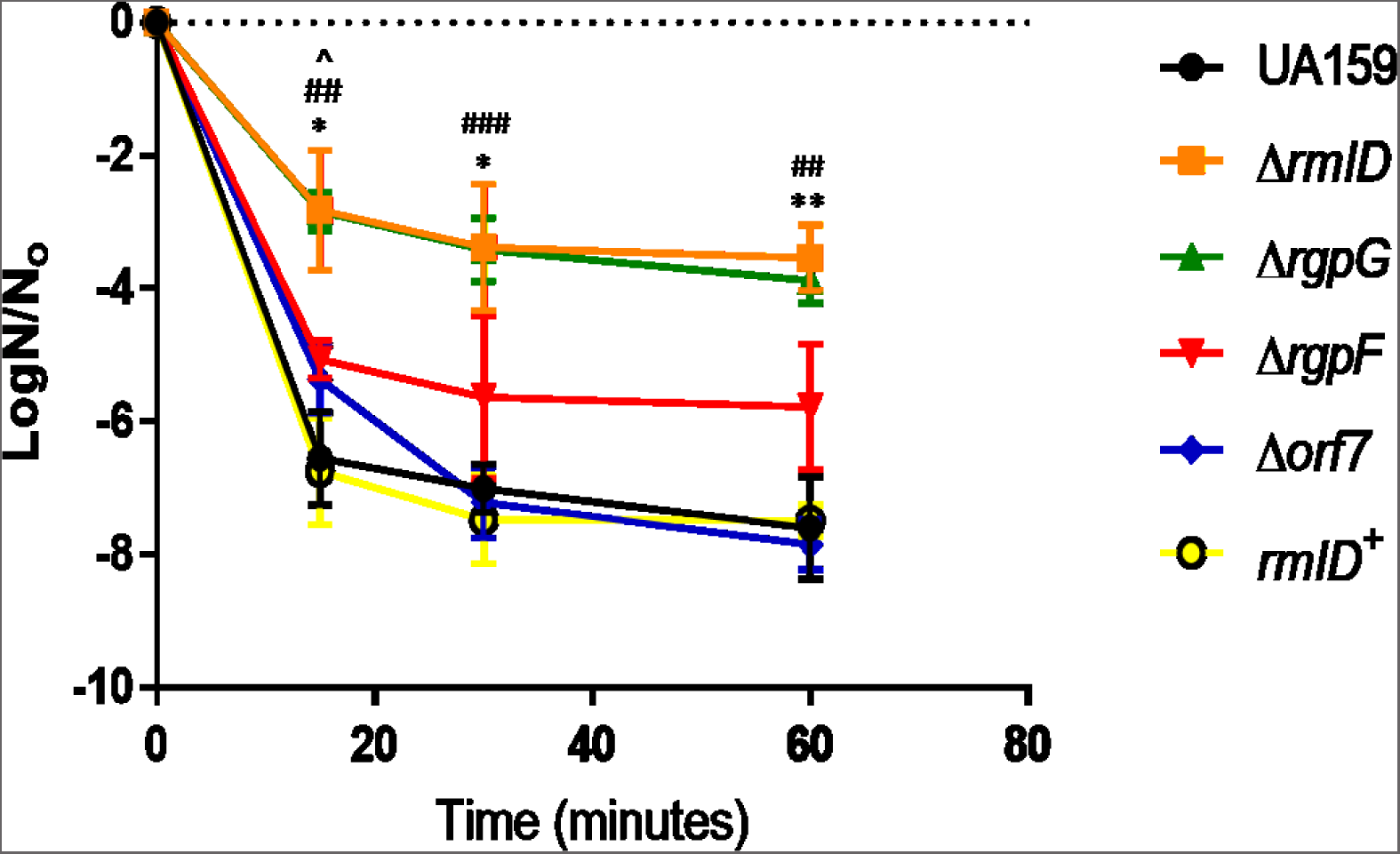
Loss of the RGP structure results in an increased resistance to lethal amounts of ferrous iron. Three independent cultures of *S. mutans* carrying deletions in *rgpF*, *rmlD*, *rgpG*, Orf7, in addition to the parent strain, UA159, and the *rmlD* complement strain (*rmlD^+^*) were grown in BHI medium to mid-log phase. Cells were exposed to 10 mM ferrous ammonium sulfate for 15, 30, and 60 minutes, serially diluted, and plated on BHI agar medium, as described in Materials and Methods. Survival was enumerated and plotted as Log (N/N_0_), where N indicates the number of surviving cells at the particular time point as a function of the number of cells at time zero (N_0_). Statistical analysis was performed using Student’s *t-*test between the indicated mutant and UA159 and is indicated by * for Δr*mlD*, # for Δ*rgpG*, and ^ for Δ*rgpF*. One symbol indicates *p* < 0.05; two symbols indicate *p* < 0.01; and three symbols indicate *p* < 0.001.

### Loss of RGP alters iron uptake

We tried to reconcile the unexpected results of the iron survival assays with previous data showing that strains with defective RGP are more susceptible to oxidative stress. We hypothesized that perhaps the RGP may be involved in iron uptake, and that the loss or truncation of the structure may lead to decreased iron uptake, allowing for reduced intracellular iron levels with fewer opportunities for damage to occur. There is precedence for this hypothesis in other Gram-positive organisms as well. In *Bacillus subtilis*, it has been shown that the negatively-charged phosphates on the wall teichoic acids (WTA) assist in the uptake of magnesium (40). The RGP in streptococci has been proposed to be orthologous to the WTA in other Gram-positive organisms, as these molecules have similar functionality, and *S. mutans* lacks a classic WTA structure (5). We hypothesized that the RGP-deficient strains have impaired iron uptake, rendering the mutant strains sensitive to streptonigrin, an antibiotic that utilizes iron as a cofactor to bind to topoisomerase and inhibit DNA synthesis (41). The transcription factor PerR has been shown to maintain iron homeostasis in *S. mutans*, and a Δ*perR* (SMU.593) mutant strain was also included as a positive control (23, 27). The test strains were grown on BHI agar medium, or BHI agar medium supplemented with 1 mM ferrous ammonium sulfate (Fe(NH_4_)_2_(SO_4_)_2_), a non-lethal concentration of iron, as demonstrated in Figure 1A. Only two of the test strains exhibited any significant changes in iron uptake, compared to UA159: the Δ*perR* strain and the Δ*rmlD* strain (Figure 3A). The reduced sensitivity of the Δ*perR* strain to streptonigrin was expected due to its role in regulation of iron homeostasis (28, 42, 43). The elevated sensitivity of the Δ*rmlD* strain to streptonigrin, however, was surprising when considered in combination with the decreased sensitivity of the strain to lethal levels of iron (Figure 2). Also, of note, the Δ*rgpG* strain, while more resistant to iron killing than UA159 (Figure 2), did not display significantly altered sensitivity to streptonigrin, compared to UA159 (Figure 3A). The contrast in results between the Δ*rmlD* and Δ*rgpG* strains was striking since both strains lack any RGP attached to their cell walls, either via loss of rhamnose production (Δ*rmlD*) or a disruption to the RGP construction machinery (Δ*rgpG*). Prior to these results, no distinction had been detected in growth or stress susceptibility between the two strains (3). Further, the results of the streptonigrin assay showed that the Δ*orf7* strain also did not differ significantly from UA159, indicating that the glycerol phosphate modification likely is not involved in iron uptake.

**Figure 3.**
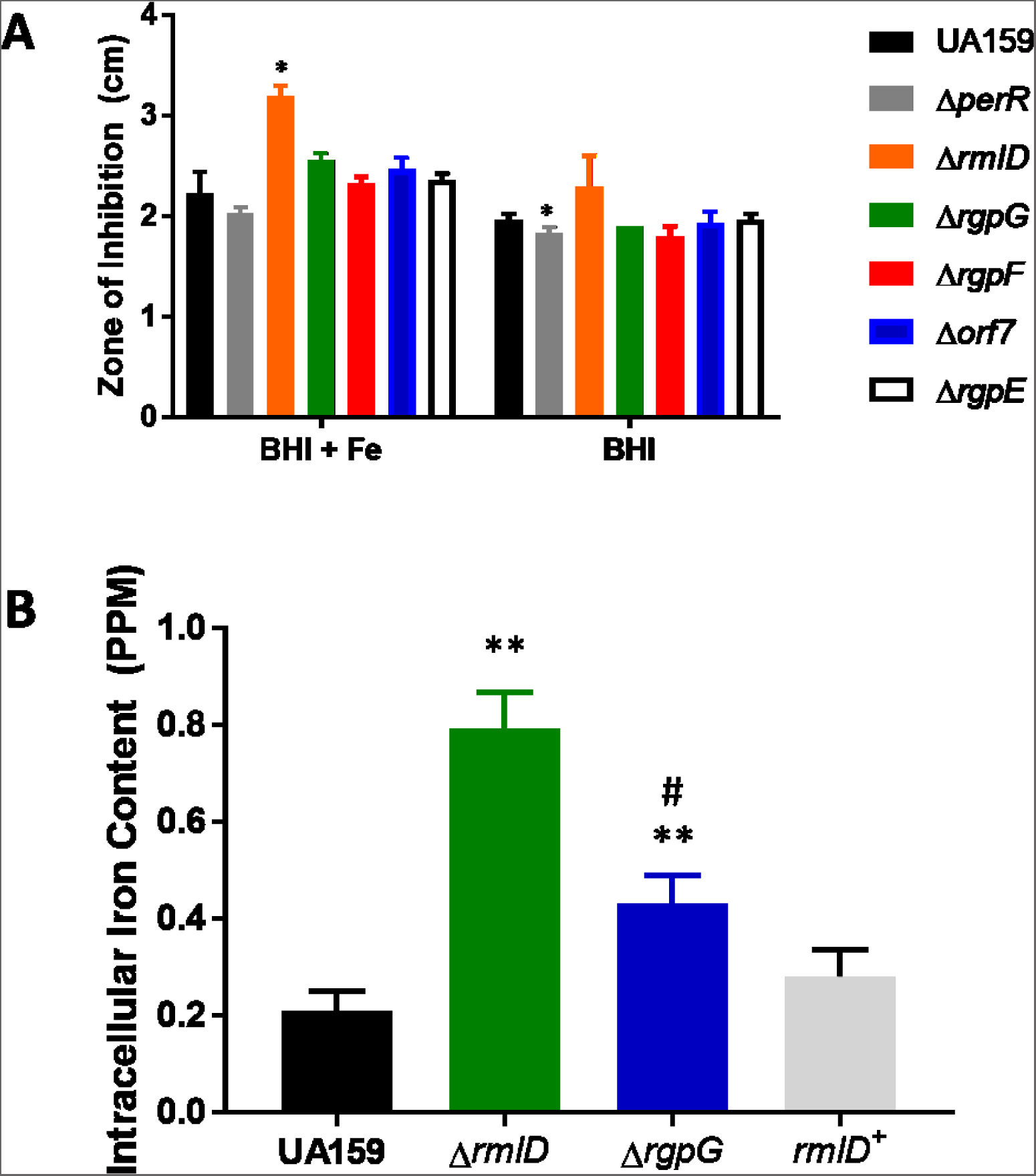
Loss of rhamnose production is tied to increased intracellular iron. Intracellular iron concentrations were measured indirectly via streptonigrin sensitivity (Panel A) or directly via ICP-OES (Panel B). **(Panel A)** Three independent cultures of *S. mutans* carrying deletions in *rgpF*, *rmlD*, *rgpG*, orf7, *rgpE*, the *rmlD* complement strain (*rmlD^+^*), the parent strain, UA159, and the Δ*perR* strain (positive control) were grown to mid-log, and an aliquot was plated on BHI agar medium with or without the addition of 1 mM ferrous ammonium sulfate. Whatman discs containing streptonigrin (5 μg/mL) were placed on the agar plate, and the plates incubated for 24 hours at 37°C. Zones of inhibition were measured to determine susceptibility to the iron-dependent antibiotic. * indicates significant difference in diameter of zones, compared to UA159, in the respective conditions (*p* < 0.05). **(Panel B)** ICP-OES was used to measure the intracellular iron content from three independent cultures of *S. mutans* UA159, and the derivative strains, Δ*rmlD*, Δ*rgpG*, and the *rmlD^+^* complement strain. Cultures were grown in BHI medium containing 1 mM ferrous ammonium sulfate, harvested, digested, and analyzed by mass spectrometry. Intracellular iron concentrations were extrapolated from a standard curve, and values were normalized to protein concentration. Statistical significance was determined between each mutant strain and UA159 using a Student’s *t-*test, **indicates significant difference in normalized iron concentration, compared to the parent strain, UA159 (*p* <0.01). ^#^indicates significant difference in intracellular iron measured in the Δ*rmlD* strain, compared to the Δ*rgpG* strain, *p* < 0.05.

Next, we wanted to directly measure intracellular levels of iron in the *S. mutans* test strains using inductively coupled plasma-optical emission spectrometry (ICP-OES) to elucidate the effects of the loss of RGP on iron uptake. There was an approx. 4-fold increase in intracellular iron levels in the Δ*rmlD* strain, compared to the parent strain, UA159 (Figure 3B), confirming our results from the streptonigrin sensitivity assays (Figure 3A). We also observed a 2-fold increase in intracellular iron in the Δ*rgpG* strain, compared to UA159 (Figure 3B). Intracellular iron levels in the *rmlD*^+^ complement strain were restored to those measured in cultures of UA159. Again, we identified distinct differences between the RGP-deficient strains, Δ*rgpG* and Δ*rmlD*. Taken together, these results are consistent with the hypothesis that increased susceptibility to oxidative stress would be linked to elevated intracellular iron content; however, the results are seemingly inconsistent with these strains being more resistant to toxic iron concentrations.

### Glucose import and proton permeability are altered in the absence of RGP

The data from the iron uptake experiments indicated a potential impairment of transport across the cell wall, so it was of interest to examine the impact of the absence of RGP on transport of other molecules. The phosphoenolpyruvate:sugar phosphotransferase (PEP-PTS) system is one of the mechanisms by which *S. mutans* can import sugars, including glucose (44). In order to further distinguish the role that *rmlD* and *rgpG* play in membrane transport, we grew cultures of these strains, along with UA159 and the *rmlD^+^* strain, prepared permeabilized cells, and measured the PEP-dependent PTS activity for glucose. The results demonstrate that the loss of the RGP structure had significant consequences on the ability of the mutant strains to import glucose via the PTS (Figure 4A). Loss of both *rmlD* and *rgpG* led to significant decreases in PTS activity (approx. a 3-fold decrease), while restoration of rhamnose production (*rmlD*^+^) produced parent strain levels of PTS activity. These results highlighted the possibility that not only iron uptake is affected by the loss of RGP/rhamnose, but that the effects of these alterations might be more global in scope.

**Figure 4.**
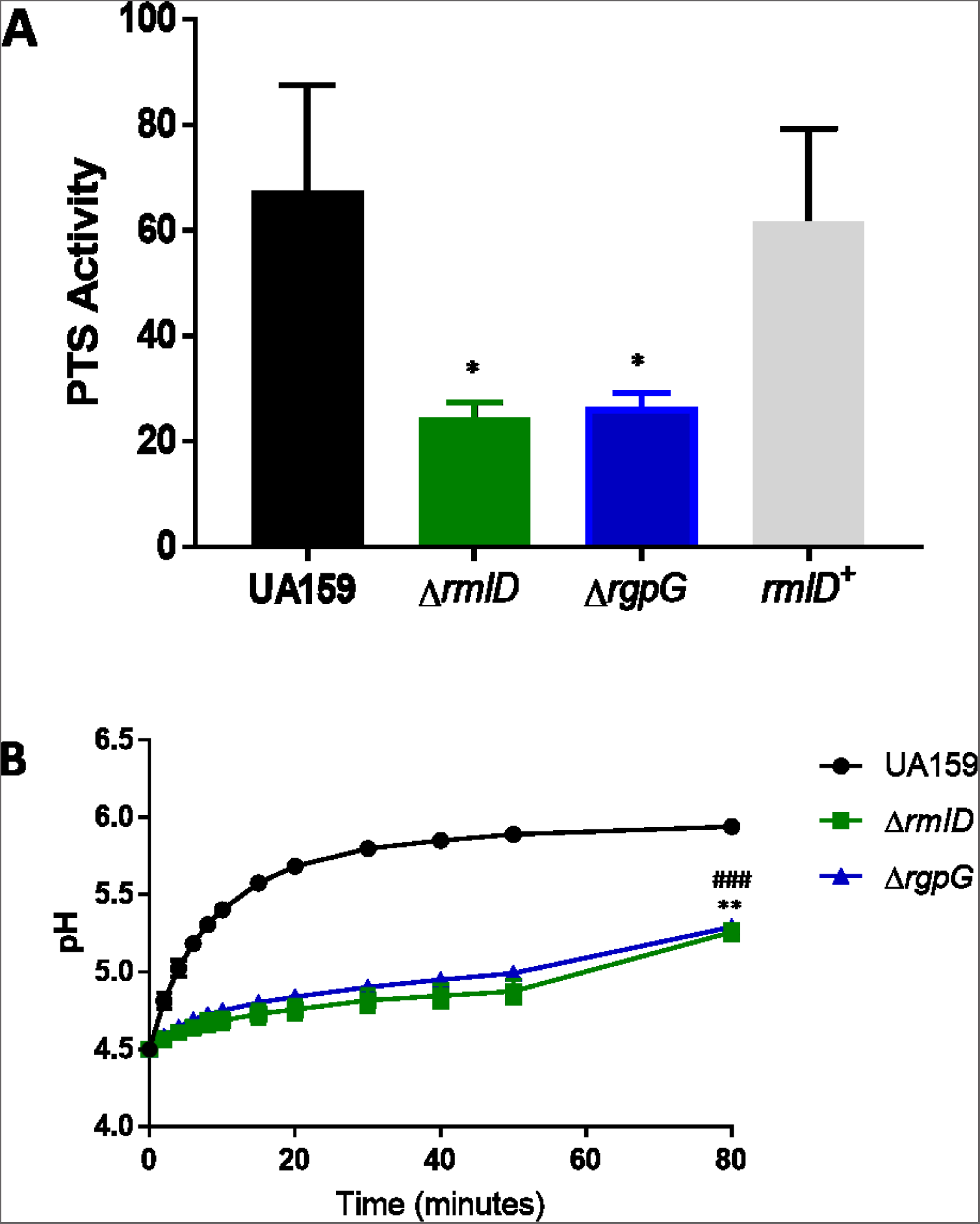
Loss of RGP disrupts membrane-associated transport. (Panel A) Glucose import via the glucose-PTS was measured in strains of *S. mutans* UA159 lacking the RGP, Δ*rmlD* and Δ*rgpG*. PTS activity is expressed as nmol pyruvate produced min^-1^ mg total protein^-1^. Statistical analysis was performed using pairwise comparison between each mutant strain and UA159, where * indicates significant decrease in PTS activity, compared to the parent strain, UA159 (*p* < 0.05); n=3. (**Panel B**). Permeability to protons was determined for strains of *S. mutans* UA159 lacking the RGP, Δ*rmlD* and Δ*rgpG*. Statistical analysis was performed using pairwise comparison between the mutant strain and UA159. **indicates a significant difference in terminal pH for the Δ*rmlD* strain, compared to the parent strain, UA159 (*p* < 0.01). ###indicates a significant difference in terminal pH for the Δ*rgpG* strain, compared to the parent strain, UA159 (*p* < 0.001); n=3 independent cultures for each test strain.

The diffusion of protons across the membrane is also an important stress tolerance mechanism in *S. mutans*. It had been previously shown that deletion of *rgpF* significantly influenced proton permeability and contributed to the sensitivity of the strain to acidic stress (4). We sought to determine if complete loss of the RGP had a similar, or more dramatic, effect on proton permeability. The loss of RGP production had severe consequences, with a drastic reduction in ΔpH over time in both the Δ*rmlD* and Δ*rgpG* strains (Figure 4B). These results provided further evidence of a diminished stress tolerance response.

### Loss of *rmlD* Causes Changes in Expression of Oxidative Stress Response Genes

In an effort to understand why the Δ*rmlD* strain exhibited increased resistance to lethal levels of iron, we decided to examine how the loss of *rmlD* might affect expression of genes known to participate in the oxidative stress response, such as *noxA*, *dpr*, and *sod* (21-24, 26, 27, 36, 45, 46). Dpr is especially relevant because of its ability to sequester iron to prevent the Fenton reaction (21–23). We measured gene expression in cultures of *S. mutans* UA159 or the Δ*rmlD* derivative grown in BHI alone, or BHI supplemented with a sublethal amount of ferrous iron (1 mM ferrous ammonium sulfate, as described above, Figure 1A). The results indicate that expression of *dpr* and *sod* increased significantly in both the UA159 and Δ*rmlD* strains in cultures grown in media supplemented with exogenous iron (Figure 5A and 5B). This was unsurprising, and confirmed that the enzymes encoded by these genes are sensitive to extracellular iron levels, and are differentially regulated to respond to those levels (21, 23, 46). There were no significant changes in expression of *noxA* in either strain grown in media supplemented with iron (Figure 5C), suggesting that *rmlD* was important only to particular aspects of the oxidative stress response. An even more intriguing finding was that expression of both *sod* and *dpr* were significantly elevated in the absence of *rmlD*: *dpr* expression increased 3-fold in cultures grown with or without ferrous iron supplementation (Figure 5A); *sod* expression increased 10-fold in cultures of the *rmlD* deletion strain grown in BHI and 8-fold in cultures grown in BHI supplemented with iron (Figure 5B). These findings could imply that the loss of *rmlD* was responsible for the altered expression levels, rather than the presence of iron, providing a plausible explanation for the increased resistance of the Δ*rmlD* strain to high concentrations of iron, compared to the parent strain.

**Figure 5.**
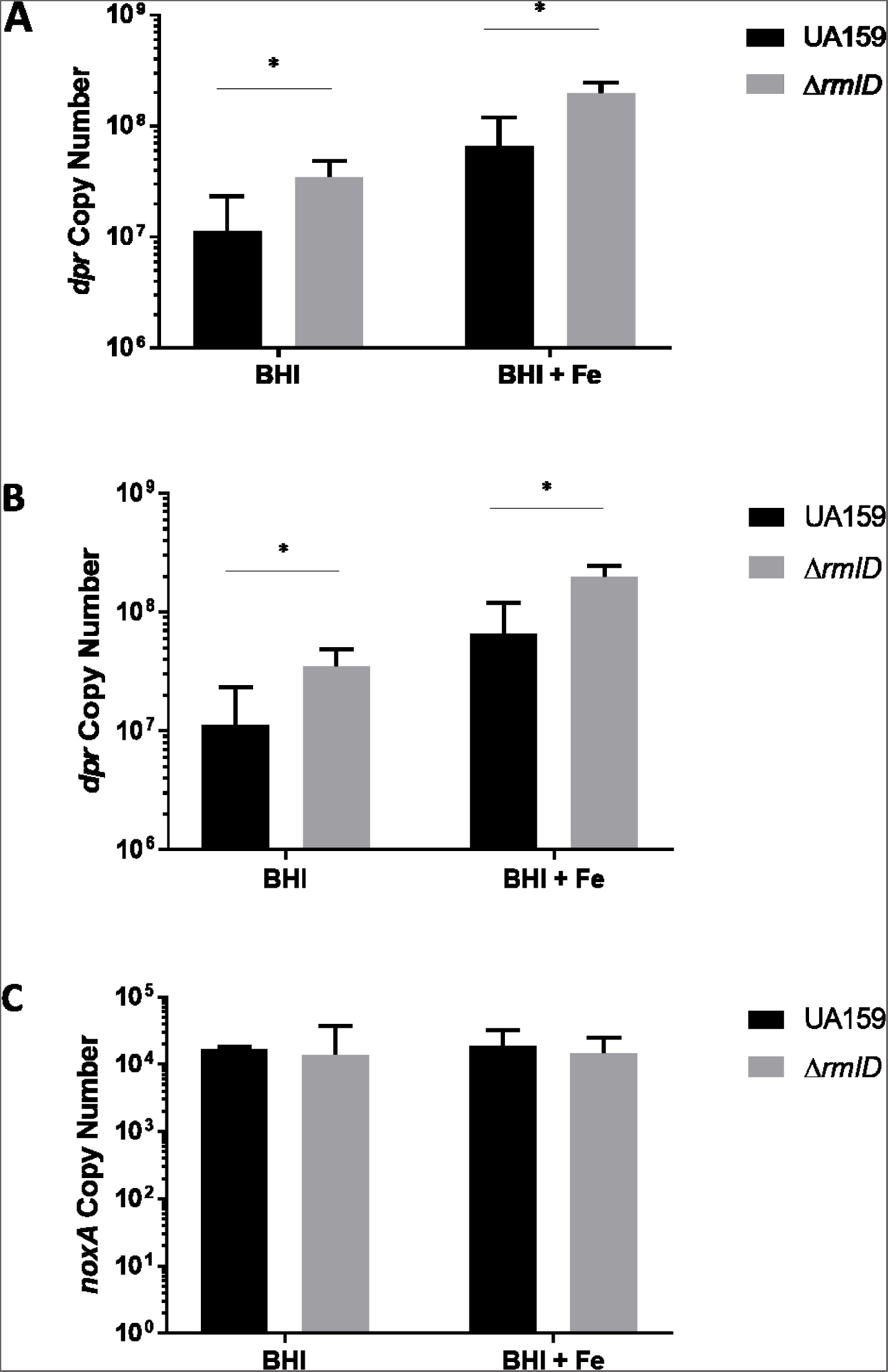
Expression of oxidative stress response genes is altered in the absence of *rmlD*. Expression of *dpr* **(Panel A)** was measured from cultures of the parent strain, UA159, and the Δ*rmlD* strain grown in BHI medium in the presence or absence of 1 mM ferrous ammonium sulfate by quantitative Real-Time PCR. RNA was isolated from three independent cultures of each strain, and qRT-PCR was performed with 3 technical replicates of each cDNA sample. Statistical significance was determined between UA159 and Δ*rmlD* in each condition using Student’s *t*-test; \**p* < 0.05. Expression of *sod* **(Panel B)** was measured from cultures of the parent strain, UA159, and the Δ*rmlD* strain grown in BHI medium in the presence or absence of 1 mM ferrous ammonium sulfate by quantitative Real-Time PCR. RNA was isolated from three independent cultures of each strain, and qRT-PCR was performed with 3 technical replicates of each cDNA sample. Statistical significance was determined between UA159 and Δ*rmlD* in each condition using Student’s *t*-test; \**p* < 0.05. Expression of *noxA* **(Panel C)** was measured from cultures of the parent strain, UA159, and the Δ*rmlD* strain grown in BHI medium in the presence or absence of 1 mM ferrous ammonium sulfate by quantitative Real-Time PCR. RNA was isolated from three independent cultures of each strain, and qRT-PCR was performed with 3 technical replicates of each cDNA sample. No statistical significance was determined between UA159 and Δ*rmlD* using Student’s *t*-test.

### *rmlD* Expression is Controlled by Regulators of the Oxidative Stress Response

The data to this point has demonstrated a role for the RGP in structural integrity of the cell and differences in expression of genes involved in the response to oxidative stress. We were next interested to learn whether the expression of the *rgp* genes could be regulated by environmental growth conditions. As *rmlD* is the first gene in the co-transcribed *rgp* operon (47, 48), we decided to focus on examining the expression of *rmlD*. We created a transcriptional fusion of the *rmlD* promoter region to a chloramphenicol acetyltransferase (CAT) reporter, to allow for measurement of *rmlD* expression as a function of CAT enzymatic activity in cultures exposed to oxidative stress conditions. Following exposure to a sublethal concentration of H_2_O_2_, we observed a significant increase in *rmlD* expression (Figure 6A). By 30 minutes post-exposure, *rmlD* expression had begun to decrease to baseline levels, likely due to the degradation of the H_2_O_2_.

**Figure 6.**
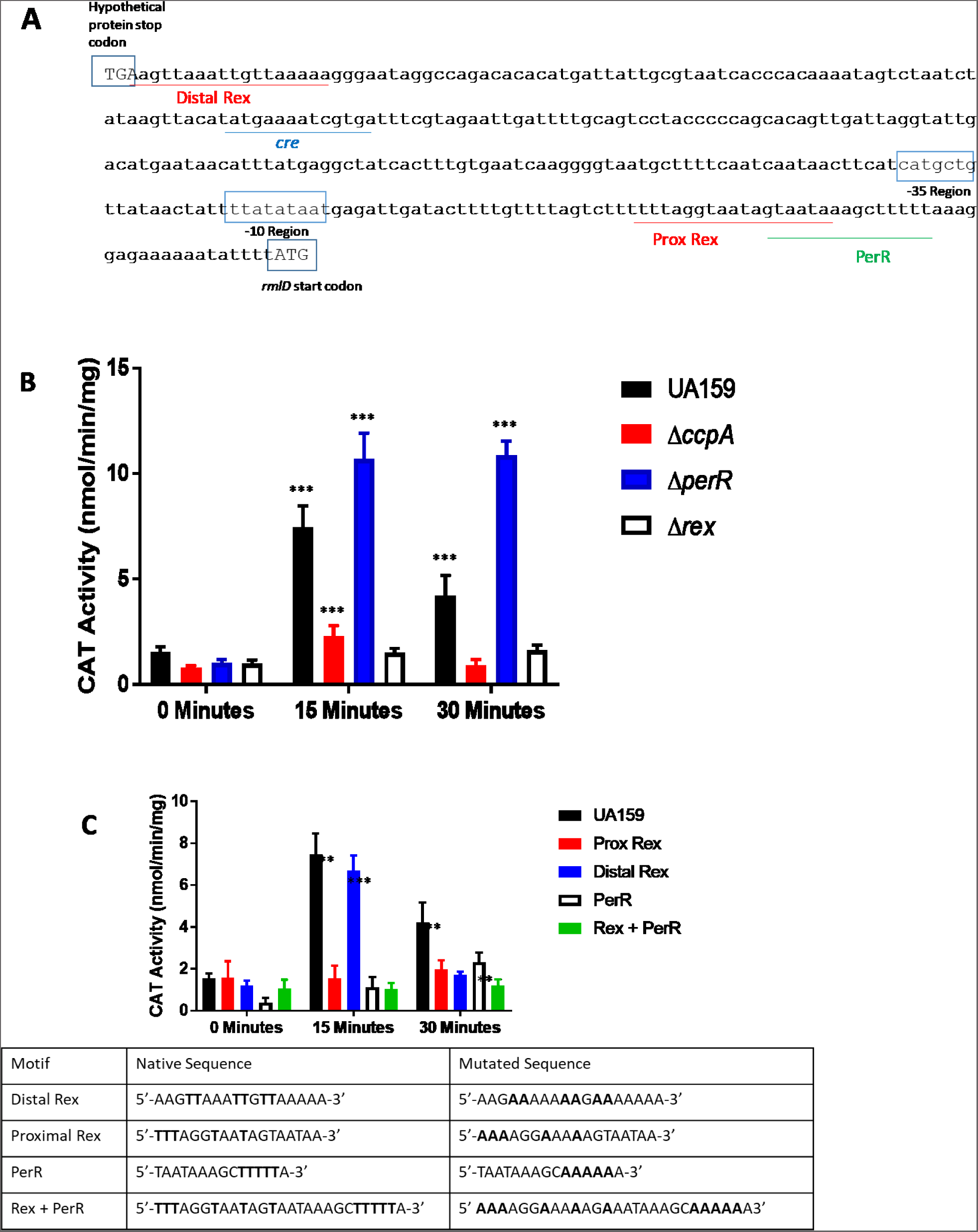
*rmlD* is regulated by the transcription factors CcpA, PerR, and Rex. (Panel A) Predicted binding motifs for CcpA (blue), PerR (green), and Rex (red) were identified in the intergenic region preceding *rmlD in silico* by Virtual Footprint. **(Panel B)** *rmlD* expression, as a function of CAT activity, was measured from extracts of three independent cultures of *S. mutans* UA159, Δ*ccpA*, Δ*perR*, and Δ*rex* background strains carrying the reporter fusion construct. Cultures were grown to mid-log phase, and then exposed to 0.5 mM H_2_O_2_ . Samples were taken 15 and 30 minutes post-addition of hydrogen peroxide and CAT activity was measured. Statistical significance was determined via pairwise comparison between each 15 and 30 minute time point, compared to time 0, for each background strain; *** indicates *p* < 0.001. **(Panel C)** Point mutations were created in the predicted binding motifs for PerR and Rex (shown in **Panel A**), as shown in the table below the figure. Expression of *rmlD*, as a function of CAT activity, was measured from *S. mutans* UA159 cultures carrying reporter fusions with these mutations. Statistical significance was determined by Student’s *t*-test between each 15 and 30 minute time point, compared to time 0, for each construct. **indicates statistical significance (*p* < 0.01); *** (*p* < 0.001).

We next were interested in identifying putative binding motifs in the *rmlD* promoter *in silico* using RegPrecise (49) that may reveal regulatory proteins involved in controlling *rmlD* expression. Interestingly, putative binding motifs for a wide variety of transcription factors were found in the intergenic space preceding *rmlD*, including regulators of DNA synthesis and global regulators (50, 51), such as CcpA, PerR, and Rex (Figure 6B). CcpA is a regulator of carbon catabolism, PerR has been shown to regulate the oxidative stress response and iron homeostasis, and Rex regulates the oxidative stress response and NAD^+^/NADH levels (23, 26, 28, 33, 34, 36, 38, 52–54). Next, H_2_O_2_-induced *rmlD* expression was examined in different strains of *S. mutans* carrying deletions in the predicted regulatory proteins CcpA, PerR, and Rex. In the Δ*ccpA* background, we saw a phenomenon similar to results from the wild-type; that is, a significant increase in *rmlD* expression at 15 minutes that returned to baseline after 30 minutes (Figure 6A). The amplitude of this response was much lower, compared to levels in the parent strain background, UA159, indicating a role for CcpA in regulation of *rmlD*, but not in response to oxidative stress. We then measured *rmlD* expression in response to H_2_O_2_ in a *perR* deletion strain, and again observed a significant increase at 15 minutes, but, interestingly, that response did not abate after 30 minutes (Figure 6A). These results suggested that perhaps PerR was controlling *rmlD* expression in response to H_2_O_2_, specifically by repressing its expression. Unlike the Δ*ccpA* (SMU.1591c) and Δ*perR* strains, there was no induction of *rmlD* expression in a Δ*rex* (SMU. 1053) strain in response to H_2_O_2_, even after 30 minutes of exposure, indicating that, similar to PerR, Rex was regulating *rmlD* expression in response to H_2_O_2_, although as an activator (Figure 6A).

CAT-derived transcription of *rmlD* was also examined in cultures grown with addition of exogenous carbohydrates or in pH-controlled conditions; however, none of those experiments yielded the significant changes in expression observed from cultures grown in the presence of H_2_O_2_ (data not shown). These results further indicated that the observed alterations in *rmlD* expression was resultant of the oxidative stress response, and not other stress response mechanisms.

While we had demonstrated that loss of the PerR and Rex regulators had distinct, but opposing, effects on the expression of *rmlD*, we wanted to determine whether the putative binding motifs found in the promoter of *rmlD* were indeed controlled by Rex and PerR, particularly because a Rex and the PerR motifs overlap in the *rmlD* promoter region. In order to answer these questions, we constructed targeted point mutations in four locations in the *rmlD* promoter: the Rex motif distal to the *rmlD* transcriptional start codon; the Rex motif proximal to the *rmlD* start codon; the PerR motif; and in a region encompassing both the proximal Rex motif and the PerR motif (Figure 6C table). Disruption of the distal Rex motif had no effect on the expression of *rmlD* when compared to expression in the wild-type background (Figure 6C). However, alteration of the proximal Rex motif resulted in no change in *rmlD* expression either 15 or 30 minutes post-addition of H_2_O_2_, analogous to results seen in the Δ*rex* strain (Figure 6C compared to Figure 6A). When the PerR motif was disrupted, *rmlD* expression was elevated at both 15 and 30 minutes after addition of H_2_O_2_ to the culture, though these increases in expression were not to the same scale as what was measured in the Δ*perR* strain (Figure 6C compared to Figure 6A). Finally, when both the proximal Rex and the PerR motifs were ablated at the same time, the result was similar to CAT activity measured from loss of the proximal Rex motif alone, indicating that binding of Rex to the *rmlD* promoter may be required for induction of *rmlD* expression in response to H_2_O_2_ (Figure 6C).

## Discussion

The rhamnose-glucose polysaccharide of *Streptococcus mutans* has increasingly been the focus of study over the last couple of decades. Earlier work focused on elucidating the roles of the genes responsible for production of the RGP, while recent work has highlighted that the presence of the RGP is important for physiological functions and for the ability of *S. mutans* to tolerate external stressors (3, 4, 6, 39). This includes both acid and oxidative stress, two common environmental stressors encountered in the oral cavity. What is unknown at this time is exactly what role the RGP plays in protecting against these stresses. Recent work investigating the function of the negatively-charged glycerol phosphate attached to the rhamnose chain has begun to shed light on this question (6). Here, we investigated the mechanistic role that RGP polymers play in providing stress tolerance to *S. mutans*, while highlighting a specifically regulated role in stress response for production of rhamnose, a fundamental building block of the RGP.

Strains of *S. mutans* with defects in RGP formation (Δ*rgpF*) or completely lacking an RGP (Δ*rmlD* and Δ*rgpG*) have been previously shown to be highly sensitive to acid and oxidative stress (3, 4, 39), while loss of the glycerol phosphate encoding gene, *orf7*, has been described to be adversely affected by the presence of zinc in culture medium (6). Therefore, it was surprising that when tested in the present study, addition of a sub-lethal concentration of ferrous iron had minimal impact on the growth rates of these *S. mutans* deletion strains, compared to growth in medium not containing the metal (Figures 1A and 1B, Table 1). This finding was observed when the cells were grown in both a microaerobic environment (55) as well as in a more oxygenated environment. A more aerobic environment (Figure 1A) would presumably favor oxidative stress conditions, especially in the presence of iron. The strains with a fully intact RGP (UA159 and *rmlD*^+^) were the only strains to display an increase in growth rate when grown in media with oxygen and iron, indicating a greater susceptibility to the stress conditions. These data would suggest that loss of the RGP was imparting some protection against iron-mediated stress.

We next quantified the susceptibility of the RGP mutant strains to a lethal concentration of iron (10 mM ferrous ammonium sulfate) via a survival assay. Again, we originally anticipated that the lack of, or disruption to, the RGP would have deleterious effects in the presence of a high dose of an oxidative stressor, but again we saw the opposite effect. The loss of RGP, produced by the deletion of either *rmlD* or *rgpG*, resulted in strains that were more resistant to the iron challenge than the wild-type, UA159 (Figure 2). This was the first evidence that loss of the RGP had a beneficial effect on the organism with regards to an oxidative stress agent. Also striking was the finding that loss of *rgpF* resulted in an intermediate level of resistance to iron-mediated killing (Figure 2). While we know that the Δ*rmlD* and Δ*rgpG* strains are devoid of cell wall rhamnose (3), it has been suggested that *S. mutans* strains lacking *rgpF* produce an altered or shortened RGP polymer with reduced cell wall rhamnose content (4). Previous studies have shown that RgpF adds the third rhamnose subunit to the rhamnose backbone of the RGP, pairing with RgpB to elongate the chain (8). While the exact structure of the RGP in a Δ*rgpF* strain has not been determined with regards to the presence of the glucose side chain and/or glycerol phosphate, our previous work has established that the growth defect exhibited by the Δ*rgpF* strain is not as severe as that observed in strains completely lacking RGP, such as Δ*rmlD* or Δ*rgpG* (3, 4). We can infer that the reduced amount of rhamnose present in the cell wall of the Δ*rgpF* strain may be enough to impart an intermediate phenotype to that of the UA159 and Δ*rgpG*/Δ*rmlD* strains with regards to iron susceptibility. Taken together, these data would suggest that it is the presence of rhamnose itself in the cell wall that plays some unknown role in coordinating iron susceptibility. It should also be noted that we saw no change in susceptibility to iron in the Δ*orf7* strain. It had previously been reported that the negatively-charged glycerol phosphate that was attached to the RGP via Orf7 played a protective role against zinc toxicity in *S. mutans* (6). We hypothesized that this negative charge might play a protective role against iron as well, but our data did not support that.

We then needed to reconcile the fact that strains typically more susceptible to oxidative stress were also more resistant to toxic iron levels. We hypothesized that strains defective in RGP had a disruption in iron uptake. Wall teichoic acid, an orthologue of the RGP in non-streptococcal Gram-positive bacteria, has been shown to play a role in uptake of magnesium (40); therefore, we wanted to determine what effect the loss of the RGP had on iron import. Streptonigrin, an antibiotic that requires iron to disrupt topoisomerase (41), has been used to assay intracellular levels of iron, such that the greater the concentration of iron within cells, the larger the zone of inhibition in a disc diffusion assay. After performing the experiments on BHI agar medium with or without 1 mM ferrous ammonium sulfate, we found that only two of the mutant strains tested (Δ*perR* and Δ*rmlD*) displayed a significant change in susceptibility to streptonigrin, compared to UA159. One of the test strains was Δ*perR*, used as a control, as the transcription factor, PerR, participates in iron homeostasis (23, 27, 28). The other streptonigrin-sensitive strain was Δ*rmlD*. This result was surprising, since it would imply that the intracellular iron content would be elevated in this strain, compared to UA159, which seemed to contradict our results from the kill assay (Figure 2). An additional unexpected result was that the difference in susceptibility between Δ*rgpG* and Δ*rmlD* was statistically significant. Both mutants lack the RGP via separate mechanisms: loss of *rmlD* prevents production of the substrate rhamnose, while deletion of *rgpG* prevents the initiation of RGP formation prior to rhamnose linkage (7). The data indicated that the test strain that had the greatest susceptibility to iron was the one that could not produce rhamnose, further suggesting that rhamnose itself may play a role in the homeostasis of intracellular iron. Results from the Δ*orf7* strain again demonstrated no significant difference in streptonigrin inhibition, compared to UA159, further indicating that the gene product plays no role in the uptake of iron.

While the streptonigrin sensitivity assay was a powerful tool, it does not directly measure intracellular iron; thus, ICP-OES was used to analyze the metal content from the various test strains. Cultures of UA159, Δ*rmlD*, and Δ*rgpG* were grown in BHI medium supplemented with 1 mM ferrous ammonium sulfate to mimic the experimental conditions used in the streptonigrin assay. The quantity of intracellular iron in the Δ*rmlD* strain was significantly increased (4-fold) over UA159, while the Δ*rgpG* strain displayed an approx. 2-fold increase in intracellular iron, compared to the parent strain. These results confirmed that loss of RGP had a significant effect on intracellular iron levels, with the highest levels found in the strain unable to produce rhamnose. Restoration of rhamnose production, in the *rmlD^+^* strain, restored intracellular iron levels to those measured in the UA159 strain, highlighting the role that the RGP plays in preserving normal iron homeostasis. These results, however, did not explain why the loss of RGP yielded an increase in intracellular iron with possible explanations including increased iron uptake, or decreased efflux of excess iron. Our results also do not provide a rationale to explain why strains with the highest intracellular iron concentrations are the most resistant to iron.

We investigated carbohydrate uptake as a means to assay transport in the absence of components of the RGP. To our knowledge, this is the first time that PTS activity has been measured in strains deficient in RGP composition. Our results demonstrate that loss of the RGP, either via deletion of *rmlD* or *rgpG* significantly diminishes PTS activity and glucose uptake, compared to the parent strain, UA159. This very likely contributes to the growth defect that is seen in RGP-deficient mutants, in addition to the replication defects (3–5, 56). The alterations in glucose import and iron uptake hint at global disruptions in nutrient uptake. Previous work from our group has established that the RGP is required for proper localization of cell wall hydrolases (5), thus, it is possible that a similar mechanism is occurring with other membrane-bound proteins. Sugar import and iron import are complex, with many different structures responsible, many with overlapping functions, and eliminating just one import mechanism does not disrupt iron uptake, indicating that any changes to import are likely to have wide-ranging causes (44, 45, 57, 58). The disruption to glucose import via the PTS gives the appearance that it may not be alterations to just iron uptake, but that there is an alteration to the uptake of a wide variety of molecules.

The observations from the assays performed thus far with the Δ*rmlD* strain have pointed towards a role for rhamnose itself in iron homeostasis, as diminishing levels of rhamnose in the RGP were associated with decreased susceptibility to iron. In *B. subtilis*, the negatively-charged WTA of the cell wall can funnel magnesium ions to ion channels for efficient uptake (40). In *S. mutans*, Orf7 adds a negatively-charged glycerol phosphate to the RGP to give the structure an overall negative charge and protect against the positively-charged zinc ion (6). We originally hypothesized that this negative charge might also play a similar role in protection against iron, but no significant changes in susceptibility to lethal levels of iron or in iron uptake were observed in the Δ*orf7* strain, compared to UA159.

Classically, iron-mediated oxidative stress is initiated through the reduction of H_2_O_2_ to the highly reactive HO• (17). Acidic conditions, or low pH, typically favor the catalysis of this reaction (59). We examined the permeability of the Δ*rmlD* strain to protons to determine whether ablation of rhamnose production altered the ability of the organism to equilibrate the ΔpH across the membrane, an important acid tolerance mechanism utilized by *S. mutans*. Our results indicate that the Δ*rmlD* strain displayed a significantly diminished ability to pump protons out of the cytosol, compared to the parent strain (Figure 4B). This data, in combination with the elevated levels of intracellular iron, would appear to favor conditions ideal for Fenton chemistry; however, the Δ*rmlD* strain was relatively tolerant of the presence of excess iron. The elevated expression of *dpr* and *sod* in the Δ*rmlD* strain may partly explain why the deletion strain is more resistant to lethal amounts of iron, while simultaneously incorporating a greater concentration of intracellular iron. As part of the acid tolerance response, *S. mutans* can prime itself, via physiological changes, to tolerate low pH (60, 61). It is possible that a similar mechanism is occurring in this case, where *dpr* and *sod* expression are elevated in the Δ*rmlD* strain, perhaps as part of the adaptation response to iron-mediated oxidative stress.

The regulators PerR and Rex are major participants in the oxidative stress response, controlling expression of *noxA*, *sod, dpr*, and *ahpCF*, among others (23, 26, 27, 34). Finding putative regulatory motifs for these regulators in the intergenic region preceding *rmlD*, and demonstrating that they affect expression of *rmlD*, suggest that RGP is a component of the oxidative stress response. Further, the overlap of the motifs for PerR and Rex highlights the control these proteins have on *rmlD* expression. Canonically, PerR is considered a repressor, while Rex has been shown to act as an activator or repressor of gene expression, dependent on the gene being regulated (30, 32, 33, 35, 37, 38). Our results support these previous findings, where *rmlD* expression was elevated in the absence of PerR, or when its binding motif was disrupted, upon exposing a culture to H_2_O_2_. The absence of Rex or alteration of its binding motif, prevented any induction of *rmlD* expression in response to H_2_O_2_. The exact mechanism by which these regulatory proteins affect expression of *rmlD* is yet to be determined. The overlapping binding motifs for these proteins, and the disparate effects of each, suggest a competitive interaction for binding to the intergenic region preceding *rmlD*. In turn, we anticipate that alterations in expression of *rmlD* can directly influence rhamnose production, and thus, the availability of the major substrate required for RGP maturation in *S. mutans*. The implications of a regulated mechanism governing cell wall polysaccharide formation may offer new insight into stress response mechanisms in Gram-positive bacteria, and these specific regulatory interactions are the subject of ongoing work.

Repression of gene expression by PerR is relieved by the H_2_O_2_-mediated oxidation of two histidine residues that causes the release of PerR from DNA (62). It is possible that PerR and Rex physically compete for binding space in the *rmlD* promoter. Due to the overlap between the two binding motifs, it is likely that only one protein can bind at a time, and the opposing effects each regulator has would support that hypothesis. One possibility for what is occurring is that PerR binds to the *rmlD* promoter under standard conditions. When the cells are exposed to ROS, PerR binding to the promoter is alleviated and the binding motif is empty. These conditions would likely favor NAD^+^ production as well, mediated by NADH oxidase, and would increase the NAD^+^/NADH ratio (20). High NAD^+^/NADH ratios mediate Rex regulation (35, 36), which would then allow Rex binding to the *rmlD* promoter and activation of transcription. Further studies will be required to interrogate the relationship between Rex and PerR in regulation of *rmlD*, although it should be noted that to this point, no studies have previously found competition between the two regulators.

In conclusion, we have demonstrated a connection between the RGP and iron homeostasis in *S. mutans*. In the presence of low concentrations of ferrous iron, complete loss of the RGP had no effect on susceptibility of the organisms to decreased growth; however, as the concentration of iron increased, the resistance to iron-mediated growth inhibition was also increased. Further, this study revealed that ablation of rhamnose production (Δ*rmlD*) had a greater impact on intracellular iron content than loss of the RGP structure alone (Δ*rgpG*). Intracellular iron concentrations in both the *rmlD* and the *rgpG* deletion strains were significantly elevated, compared to the parent strain, UA159. In addition to alterations in iron homeostasis, loss of the RGP structure also resulted in impaired glucose uptake and reduced proton permeability of the membrane, compared to UA159. The increased resistance to iron observed in the Δ*rmlD* strain may be partially explained by increased expression of *dpr* and *sod*, two genes encoding products responsible for protection against oxidative stress. Finally, our data demonstrates that expression of *rmlD* is regulated by the oxidative stress response regulators Rex and PerR, with the latter is also involved in iron homeostasis, highlighting a role for the RGP in protection against oxidative stress and in facilitating iron uptake.

## Materials and Methods

### Bacterial Strains and Growth Conditions

*Streptococcus mutans* UA159 was used as the wild-type strain in all experiments conducted (47). All deletion strains were created by replacement of the open reading frame with an erythromycin resistance cassette, as previously described (3, 4, 39). The *rmlD*^+^ strain was created by integrating a copy of *rmlD* into the *gtfA* locus, followed by deletion of *rmlD* in the native locus, as previously described (3). All cultures were grown in brain heart infusion medium (BHI; BD/Difco, Franklin Lakes, NJ) unless otherwise noted.

### Bacterial Growth Curves

Growth rates were measured for all strains using a Bioscreen C plate reader (Growth Curves USA, Piscataway, NJ) as previously described (3). Briefly, overnight cultures were diluted 1:10 into BHI and allowed to grow at 37°C in a 5% (vol/vol) CO_2_–95% air atmosphere to mid-log phase. From the culture, a 10μL aliquot was diluted into fresh BHI which was supplemented with 1 mM ferrous ammonium sulfate (Fe(NH_4_)_2_(SO_4_)_2_). These strains were then allowed to grow at 37°C aerobically and absorbance readings (OD_600_) were taken every 15 minutes over a period of 24 hours. Prior to each reading, cultures were mixed with a 10 second period of shaking at medium amplitude. For experimentation under microaerobic conditions, mineral oil was added on top of growth medium to create an oxygen-buffering overlay. The experiments were performed using 3 independent cultures, with ten technical replicates each.

### Iron Susceptibility Assay

Overnight cultures of *S. mutans* deletion strains, Δ*rmlD*, Δ*rgpF*, Δ*rgpG*, Δ*orf7*, the *rmlD*^+^ complement strain, and the parent strain, UA159, were diluted 1:20 into 30mL of fresh BHI. Cells were allowed to grow to mid-log and then harvested. The pellet was then resuspended in 9 mL of fresh BHI. Prior to addition of iron, 0.1 mL of the resuspended *S. mutans* cultures were removed, serially diluted to 10^-8^, and plated on BHI agar medium. This was time zero. Then, ferrous ammonium sulfate was added to the culture, to a final concentration of 10 mM. The culture was then incubated in a 37°C water bath for the duration of the experiment. At 15, 30, and 60 minutes post-addition of the ferrous ammonium sulfate, 0.1 mL of the culture was removed, serially diluted, and plated, as mentioned above. Plates were then incubated at 37°C in a 5% (vol/vol) CO_2_–95% air atmosphere for 48 hours before colonies were enumerated and viable colony forming units were determined. The experiment was performed in triplicate, using 3 independent cultures of each strain.

### Streptonigrin Sensitivity Assay

Sensitivity of *S. mutans* strains to streptonigrin (Sigma-Aldrich, St. Louis, MO) was tested via disc diffusion assays. Cultures were grown to mid-log phase and 100 μL was spread on BHI agar plates or BHI agar supplemented with 1 mM ferrous ammonium sulfate. Streptonigrin, 2 μM (in 99% EtOH) or 99% EtOH (vehicle control), was directly pipetted onto Whatman filter paper discs (6mm diameter) (Cytiva, Marlborough, MA). Three discs, as technical replicates, were placed on each agar plate, and each strain was tested in triplicate from 3 independent cultures. The plates were allowed to incubate for 24 hours at 37°C in a 5% (vol/vol) CO_2_–95% air atmosphere. Following incubation, the diameter of the zone of inhibition was measured for each replicate, the measurements averaged, and standard deviation calculated.

### Intracellular Iron Quantification

Intracellular iron content of *S. mutans* strains Δ*rmlD*, Δ*rgpG*, the *rmlD^+^* complement strain, and the parent strain, UA159, was measured using inductively coupled plasma-optical emission spectrometry (ICP-OES) at the University of Rochester Medical Center Environmental Analysis Facility. The assay was completed as previously described (45, 63). Briefly, cultures of *S. mutans* were grown in BHI medium containing 1 mM ferrous ammonium sulfate. After reaching mid-log phase, the cultures were harvested, washed twice in ice-cold PBS containing 0.5 mM EDTA, then washed once with PBS alone. The cell pellets were resuspended in 4 mL of 35% TraceMetal^TM^ Grade nitric acid (Fisher Scientific), placed in a vial (HDPE) (VWR, Radnor, PA), and incubated at 90°C for 1 hour. Following digestion, samples were diluted 1:10 in reagent-grade H_2_O and iron concentrations were quantified from the diluted samples using a Perkin Elmer Avio 200 mass spectrometer (Perkin Elmer, Waltham, MA) and compared to a standard curve. Iron concentrations were then normalized to the protein content in each sample using the bicinchoninic acid (BCA) assay (Sigma Aldrich, St. Louis, MO). Quantification was performed from 3 independent cultures for each strain.

### Phosphoenolpyruvate:sugar Phosphotransferase System (PEP-PTS) Assay

To measure phosphoenolpyruvate-dependent glucose uptake, a PTS assay was performed as previously described (36, 44). Briefly, a 50 mL overnight culture was grown in tryptone-yeast extract (TY) media + 1% (w/v) glucose (TYG). Cells were harvested and washed twice with 0.1M sodium-potassium phosphate buffer (pH 7.2) containing 5 mM MgCl_2_. The pellet was resuspended in 5 mL of the same buffer, and placed on ice. Toluene:acetone (1:9) was added to the cell suspensions, the tubes were vortexed for two minutes, placed on ice for two minutes and then vortexed again for two minutes. A master mix for the assay containing 0.1 mM NADH, 10 mM NaF, 10 mM glucose, 10 U of lactic acid dehydrogenase, and 0.16 M sodium-potassium phosphate buffer was made and pipetted into each of 4 cuvettes. The cuvettes were warmed to 37°C for 5 minutes, then 50 μL of cell suspension was added to each cuvette and the mixture blanked at OD_340_. The reaction was initiated by addition of 10 μL of 0.5 M PEP and optical density readings were taken every ten seconds for two minutes. The data was normalized to protein concentration, as determined by BCA assay. Results are presented as PTS activity over time, where PTS activity represents nmol pyruvate produced min^-1^ mg total protein^-1^. The experiments were performed in triplicate, using 3 independent cultures.

### Proton Permeability Assay

The ability of *S. mutans* strains to allow protons across their membrane was measured as previously described (4, 64). Briefly, 200 mL cultures of *S. mutans* UA159, or its derivatives, Δ*rmlD* and Δ*rgpG*, were grown overnight and the OD_700_ was measured in the morning. The cells were harvested, washed with 5 mM MgCl_2_, and resuspended in 20 mM phosphate buffer (pH 7.2) containing 1 mM MgCl_2_/50 mM KCl to a final cell concentration of 5 mg cell dry weight per mL. The samples were then starved for 120 minutes at 37°C. After starvation, cells were once again harvested and resuspended in 50 mM KCl/1 mM MgCl_2_ to a final cell concentration of 20 mg cell dry weight per mL. The pH of this solution was rapidly adjusted to 4.5 (designated 0 minutes) and pH readings were taken over the next 50 minutes. At 50 minutes, butanol was added to a final concentration of 10% (vol/vol) and a final pH reading was taken at 80 minutes.

### Real-time Quantitative PCR

Briefly, four independent cultures of the parent strain, *S. mutans* UA159, and the Δ*rmlD* strain were grown to mid-log phase in BHI medium, and harvested. RNA was extracted using previously described methods (36). RNA (500 ng) from each strain was amplified with random primers to synthesize cDNA with the High-Capacity cDNA Reverse Transcription Kit (Life Technologies, Carlsbad, CA). Primer pairs listed in Table S1 were used to amplify cDNA to measure expression of the *S. mutans* genes *sod, dpr,* and *noxA* by qRT-PCR using PowerSYBR Green master mix (Life Technologies) in a StepOnePlus Real Time PCR System (Applied Biosystems, Foster City, CA). The concentration of the DNA templates were used to determine copy number and serial dilutions were performed to create a standard curve for each target gene. The mRNA copy number was quantified based on a standard curve, as previously described (36).

### Chloramphenicol Acetyltransferase Assays

The intergenic region preceding *rmlD* was PCR amplified from UA159 genomic DNA with primer pair rmlDSacI-PROM-FOR and rmlDBamHI-PROM-REV. The amplicon was gel isolated, sequentially digested with *Sac*I and *Bam*HI (New England Biolabs, Ipswich, MA), and purified by gel electrophoresis. The purified *rmlD* promoter region was cloned into the promoterless chloramphenicol acetyltransferase reporter plasmid pJL84, digested with *Bam*HI and *Sac*I (65). The ligation mixture was transformed into *E. coli* DH10B and colonies were selected on LB medium containing kanamycin (50 μg mL^-1^). Colony PCR using the primer pair rmlDSacI-PROM-FOR and CATJL (Table S1) was used to confirm constructs and a sequence-verified clone was designated pJLrmlD. Purified plasmid DNA was then transformed into *S. mutans* UA159 to generate a single-copy, integrated reporter fusion. Transformants were selected on BHI agar medium containing 1 mg mL^-1^ kanamycin and screened by colony PCR using the primer pair rmlDSacI-PROM-FOR and CATJL to verify proper chromosomal integration.

Three independent cultures of *S. mutans* strains UA159, Δ*ccpA* (39, 66), Δ*perR* (39), and Δ*rex* (36, 39), carrying pJLrmlD were grown to mid-logarithmic phase in BHI medium. An aliquot of the culture was removed at time 0, then hydrogen peroxide was added to the remaining culture to a final concentration of 0.5 mM. Aliquots (50 mL) were removed from the cultures 15- and 30-minutes post-addition of H_2_O_2_, and harvested (2272 x *g*, 4⁰C). The cell pellets were washed once with 10 mM Tris-Cl (pH 7.8) and centrifuged again as above. Measurement of *rmlD* promoter activity as a function of CAT activity was performed as previously described (26). Briefly, cell pellets were resuspended in 10 mM Tris-Cl (pH 7.8) and transferred to a screw-capped tube containing 0.1 mm glass beads (BioSpec Products, Bartlesville, OK). Cells were disrupted in a BeadBeater 8 (BioSpec Products) for two 30 second pulses, with an incubation on ice in between. Cleared lysates were obtained after centrifugation at 15,871 x *g* at 4⁰C. Each CAT assay reaction mixture consisted of 50 µL cell lysate; 100 mM Tris-Cl, pH 7.8; 0.1 mM acetyl-coenzyme A (acetyl-CoA); and 0.4 mg ml^-1^ 5,5-dithio-bis (2-nitrobenzoic acid) (DTNB) in a total volume of 1 ml. Reactions were initiated by addition of 0.1 mM chloramphenicol. Optical density measurements at 412 nm were monitored over 3 min. The reaction rate and total protein concentration (determined by the method of Bradford with the BioRad Protein Assay reagent; BioRad, Hercules, CA) were used to determine CAT activity. Results are presented as nmol of chloramphenicol acetylated min^-1^ mg total protein^-1^.

### Site-directed Mutagenesis

Targeted mutations were created in the Rex and PerR motifs located in the *rmlD* promoter region of pJLrmlD, as described in the table in Figure 6C. Synthetic DNA constructs of the 331bp *rmlD* promoter region carrying point mutations in the distal Rex, proximal Rex, PerR, or proximal Rex + PerR motifs were constructed by Genscript (Piscataway, NJ) in the *Eco*RV site of pUC57. Plasmid DNAs for each of the 4 constructs were sequentially digested with *Sac*I and *Bam*HI (New England Biolabs, Ipswich, MA), purified by gel electrophoresis, and cloned into similarly digested pJL84, as described above for the native *rmlD* construct. Purified plasmid DNA from each reporter fusion construct was then transformed into *S. mutans* UA159 to generate a single-copy, integrated reporter fusion. Transformants were selected on BHI agar medium containing 1 mg mL^-1^ kanamycin and screened by colony PCR using the primer pair rmlDSacI-PROM-FOR and CATJL to verify proper chromosomal integration.

### Statistical Analysis

Statistical significance was determined by pairwise comparison using Student’s *t* test.

## Acknowledgments

We thank Matthew D. Rand and Thomas Scrimale (University of Rochester Medical Center, Rochester, NY) for their assistance with the ICP-OES experimentation and analysis.

This study was supported by NIH/NIDCR DE-13683 and DE-17425 (to R.G.Q.) and the NIH/NIDCR Training Program in Oral Sciences DE021985 (to A.P.B. and C.J.K.).

