## Supplemental Figures for "The *Streptococcus mutans* Rhamnose-glucose Polysaccharide Plays an Important Role in Oxidative Stress Resistance and Iron Homeostasis"

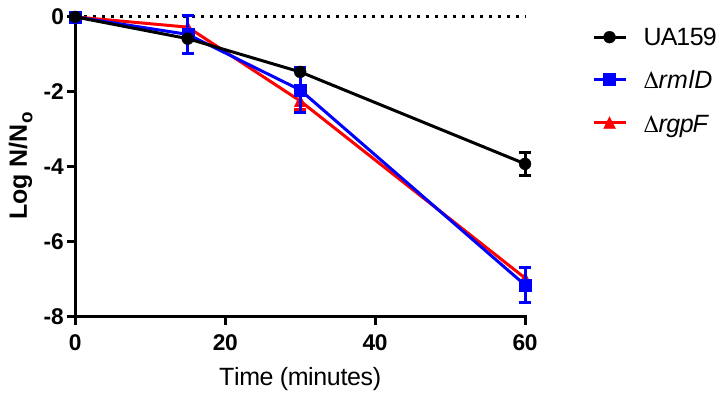

***

**Figure S1. Loss or alteration of the RGP increases susceptibility of *S. mutans* to H_2_O_2_.** Three independent cultures of *S. mutans* UA159, Δ*rmlD*, and Δ*rgpF* were grown to mid-log and harvested. Cell pellets were resuspended in BHI medium, serially diluted, and plated (time 0). Hydrogen peroxide (16.3 mM H_2_O_2_) was added to the cell suspension, and aliquots were removed after 15, 30, or 60 minutes, serially diluted, and plated on BHI agar medium. CFU were enumerated, and survival is represented as Log (N/N_0_). Statistical significance was determined by Student’s *t*-test for the pairwise comparison between either the Δ*rmlD* or the Δ*rgpF* strain, compared to UA159, at each time point; *** indicates *p* < 0.001.

**Table S1:** List of oligonucleotide primers used in this study.

| Primer | Sequence (5’-3’) | Source |
| --- | --- | --- |
| rmlDSacI-PROM-FOR | CGTTGAGAGCTCAGTTAAATTGTTAAAAAGGG | This study |
| rmlDBamHI-PROM-REV | GATTAAAATCATGGATCCTATTTTTTCTCCTTTAAAAAGC | This study |
| CAT JL | TTTCTGTGGTTATACTAAAAGTC | (65) |
| dpr Fwd | GAAGAAACAGTTGGCACATGGG | (26) |
| dpr Rev | TTCCGTTTGAGCTGCTGTAAAG | (26) |
| sod Fwd | AGCACTTGATGTCTGGGAACAC | (26) |
| sod Rev | CGGCATAAAGACGAGCAACAG | (26) |
| noxA Fwd | GGGTTGTGGAATGGCACTTTGG | (26) |
| noxA Rev | CAATGGCTGTCACTGGCGATTC | (26) |
