## Supplemental Methods for "The *Streptococcus mutans* Rhamnose-glucose Polysaccharide Plays an Important Role in Oxidative Stress Resistance and Iron Homeostasis"

**Supplemental Materials for Bischer et al., 2021.**

**Supplemental Materials and Methods**

**Hydrogen Peroxide Susceptibility Assay**

Overnight cultures of *S. mutans* strains, Δ*rmlD*, Δ*rgpF*, and UA159 were diluted 1:20 into 30mL of fresh BHI. Cells were allowed to grow to mid-log and then harvested. The pellet was then resuspended in 9 mL of fresh BHI. Prior to addition of H_2_O_2_, 0.1 mL of the resuspended *S. mutans* cultures were removed, serially diluted to 10^-8^, and plated on BHI agar medium. This was time zero. Then, H_2_O_2_ was added to the culture to a final concentration of 16.3 mM. The culture was then incubated in a 37°C water bath for the duration of the experiment. At 15, 30, and 60 minutes post-addition of the H_2_O_2_, 0.1 mL of the culture was removed, serially diluted, and plated, as mentioned above. Plates were then incubated at 37°C in a 5% (vol/vol) CO_2_–95% air atmosphere for 48 hours before colonies were enumerated and viable colony forming units were determined. The experiment was performed in triplicate, using 3 independent cultures of each strain.
